# Rgg_144_/SHP_144_ - controlled Streptolancidin D mediates intra-species competition in *Streptococcus pneumoniae* with cumulative effect from other bacteriocins and fratricide

**DOI:** 10.1101/2025.09.26.678837

**Authors:** Carina Valente, Sofia Dias, Ozcan Gazioglu, Ana R. Cruz, Adriano O. Henriques, Hasan Yesilkaya, N. Luisa Hiller, Raquel Sá-Leão

## Abstract

*Streptococcus pneumoniae* is a major colonizer of the human nasopharynx, where inter- and intra-strain competition plays a critical role in shaping population structure and influencing vaccine outcomes. Bacteriocins are key mediators of intra-species competition, yet many of their functions and regulatory mechanisms remain poorly understood. Here, we identify and characterize streptolancidin D, a previously uncharacterized bacteriocin encoded by the *sldA-T* locus and demonstrate its contribution to pneumococcal competition. Using isogenic streptolancidin-producing and non-producing variants of a naturally colonizing strain, we show that *sldA-T* contributes to inhibition of competitor strains in *in vitro* biofilms and during murine co-colonization. Importantly, streptolancidin D also inhibited a subset of genetically diverse pneumococcal isolates representing multiple serotypes, whereas non-producing variants showed no activity. This indicates that its effect is broad and not restricted to isogenic interactions. Genomic analysis of over 7,500 pneumococcal genomes revealed that *sldA-T* is present in ∼12% of isolates, with lineage-associated distribution patterns, and is consistently encoded downstream of the Rgg_144_/SHP_144_ quorum sensing system. We further demonstrate that *sldA-T* is regulated by this system, with *sldA-T* promoter activity abolished in a SHP-deficient background and partially restored by exogenous peptide stimulation. Finally, we show that streptolancidin D acts in concert with other bacteriocin systems and competence-mediated fratricide, highlighting a multifactorial antimicrobial strategy that enhances pneumococcal competitiveness. Overall, our findings identify a quorum sensing-regulated bacteriocin that contributes to pneumococcal competition and helps shape population dynamics.

**IMPORTANCE:** Bacteriocins are central to bacterial competition and niche occupation, particularly in structured environments like the human nasopharynx. While several pneumococcal bacteriocins have been characterized, the functions of many remain unknown, limiting our understanding of how these systems shape strain fitness and population dynamics. We characterize streptolancidin D, a bacteriocin that enhances intraspecies competitiveness *in vitro* and *in vivo* and contributes to inhibition of genetically diverse pneumococcal strains. We demonstrate that its expression is tightly regulated by the conserved Rgg_144_/SHP_144_ quorum sensing system and that the locus is distributed and shows synteny across multiple pneumococcal lineages. Our findings reveal that streptolancidin D operates within a broader network of bacteriocins and competence-associated mechanisms that collectively shape competitive interactions. By integrating genomic, functional, and regulatory analyses, this work expands the known repertoire of pneumococcal antimicrobial systems and provides new insights into the mechanisms underpinning competition and population structure in *S. pneumoniae*.

## INTRODUCTION

*Streptococcus pneumoniae* (pneumococcus) is a major human pathogen responsible for considerable global morbidity and mortality. Invasive disease, however, is relatively rare compared to the high frequency of asymptomatic nasopharyngeal colonization, especially in children (1). Individuals are often colonized by multiple pneumococcal strains, either simultaneously or sequentially (2–4), but very little is known about the mechanisms by which co-colonizing strains interact.

In the nasopharynx, pneumococci form biofilms - structured communities embedded in extracellular matrix - on mucosal surfaces, which enhance persistence and protect against host defenses (5). Within this environment, strains must compete for space and nutrients, leading to a dynamic network of intra- and inter-species interactions.

Bacteriocins, ribosomally synthesized antimicrobial peptides, are key mediators of intra-species competition by inhibiting or killing neighboring strains that lack the cognate immunity genes (6–11). Pneumococcal genomes encode multiple bacteriocin loci, ranging from ubiquitous systems like the pneumocins (or bacteriocin-like peptides, Blp) and competence-induced bacteriocins (CibAB), to more variable loci such as streptolancidins, streptococcins, and pneumolancidin K (PldK) (8, 12, 13). Despite their prevalence, the regulatory mechanisms and functional roles of most of these loci remain largely unexplored. Exceptions include Blp, CibAB, streptococcin B (Scb), and PldK, which have been functionally characterized and shown to mediate intra-species antagonism (6, 8, 9, 14).

The Blp system is regulated by the BlpC pheromone through the BlpRH two-component system, leading to expression of a polymorphic bacteriocin-immunity region (BIR) (6, 11). The CibAB system is activated by the competence-stimulating peptide (CSP) via the ComDE two-component system and functions as part of competence-mediated fratricide, a cell-to-cell contact-dependent mechanism in which competent cells lyse non-competent neighboring siblings through effectors such as the murein hydrolase CbpD and the competence-induced bacteriocins CibA and CibB. This mechanism, thereby promotes DNA release, horizontal gene transfer, and competitive fitness during colonization (7, 14, 17, 18). Although BlpC and CSP activate different systems, functional promiscuity of their ABC transporters allows cross-regulation, linking *blp* expression to the competence regulon (15, 16). The recently described *scb* locus, encodes a lactococcin 972-like bacteriocin. Expression of *scbABC* is regulated by ComE during competence and, in some strains, it can also be activated via a ComR-type quorum sensing system in response to *comX*-inducing peptides (XIP). Scb activity enhances fitness by targeting *scb*-negative or non-competent strains, functioning both in fratricide and inter-strain competition (9). Together, these systems underscore the close coordination between bacteriocin production, competence, and competitive interactions in pneumococci. Quorum sensing systems of the Rgg/SHP family have emerged as key regulators of cell-cell communication and may play a broader role in controlling antimicrobial peptide production than previously appreciated (reviewed in (19)). In fact, Javan et al. have proposed that the region downstream of the Rgg_144_/SHP_144_ constitutes a bacteriocin cluster hotspot, where bacteriocin-associated gene clusters frequently insert into the pneumococcal genome (12). This system consists of Rgg_144_, a transcriptional regulator, and SHP_144_, a short hydrophobic signaling peptide. SHP_144_ is secreted, accumulates with increasing cell density, and is internalized via the Opp oligopeptide permease; it then binds Rgg_144_ and triggers DNA-binding activity (20). In *S. pneumoniae* Rgg_144_/SHP_144_ controls the transcription of genes associated with biofilm formation and virulence (21, 22).

In this study, we explored the mechanism of known inhibitory activity of a naturally colonizing 19F pneumococcal strain towards other pneumococci in *in vitro* biofilms (23). We hypothesized that secreted bacteriocins were responsible for the observed inhibitory activity. We identified bacteriocin streptolancidin D, encoded by the *sldA-T* locus, as mediator of inhibition. Given the lack of functional studies on this locus, we characterized the role of the *sldA-T* locus in mediating inhibition in vitro and *in vivo*, mapped its distribution across pneumococcal lineages, and examined its regulation by the Rgg_144_/SHP_144_ system. We further show its interplay with other bacteriocin loci and competence-mediated fratricide.

## RESULTS

### Secreted peptides are involved in the inhibitory activity of strain 19F

We have previously identified a pair of naturally colonizing pneumococci with a strong negative interaction in biofilm in which a strain 1990-19F, herein called 19F, strongly inhibits the growth of the non-encapsulated 7031-NT strain, herein called NT, and other pneumococcal strains, including the laboratory strain D39 (23).

We hypothesized that this inhibitory effect could be attributable to a secreted molecule. To test this hypothesis, cell-free supernatants (CFS) of early stationary-phase planktonic cultures of strain 19F were obtained from peptide free chemically defined medium (CDM) and added to cultures of NT strain at the time of biofilm inoculation to a final concentration of 20% (v/v). The CFS of 19F had significant inhibitory activity towards the NT strain biofilm. This activity was significantly reduced upon treatment of the CFS with a broad-spectrum protease but was maintained upon treatment at 65°C. The reduced cell counts of NT biofilms were not caused by medium dilution resulting from addition of spent medium, since adding 20% water (v/v) resulted in cell counts comparable to the control conditions. Of note, protease treatment did not completely abolish the inhibitory activity of the CFS of the 19F strain towards NT biofilms (**Figure 1**).

**Figure 1.**
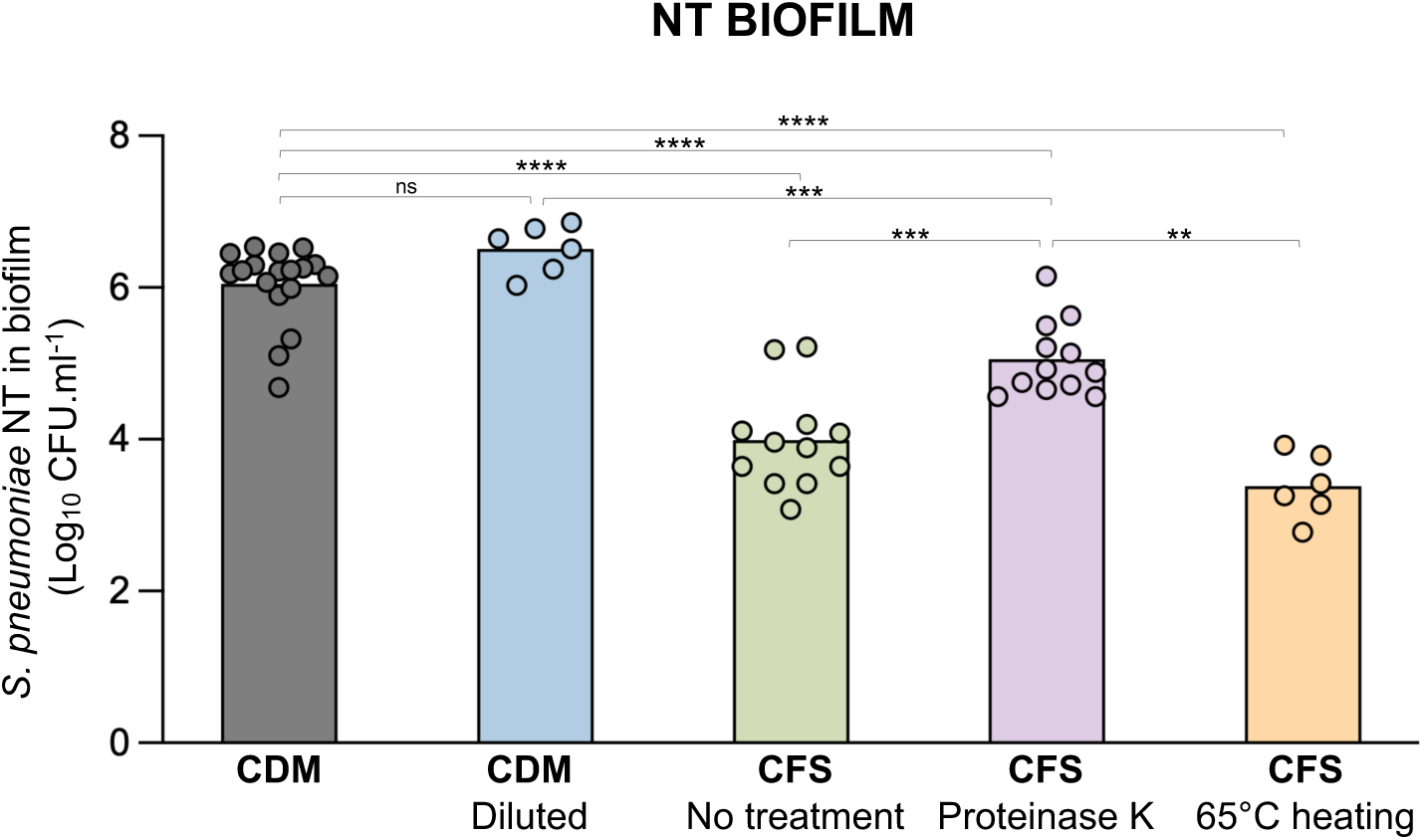
The inhibition of strain NT is mediated by a proteinaceous molecule secreted by strain 19F. Biofilms were grown in CDM for 24h and 20% (v/v) of cell-free supernatant (CFS) obtained from 19F cultures were added at time of inoculation. CDM, no addition of supernatant; Diluted, CDM diluted with 20% (v/v) water; No treatment, addition of non-treated supernatant; Proteinase K, addition of supernatant treated with 1mg.mL^-1^ proteinase K for 1h at 37°C, followed by heat inactivation; Heat-treated supernatant, addition of supernatant heated to 65°C for 20min; Events.mL^-1^ were calculated based on cell counts by flow cytometry of resuspended NT biofilms. Plot represents the results of at least three independent experiments, each with intra-experiment duplicates. Black bars indicate geometric means. ns, non-significant; *, *p*-value <0.05; **, *p*-value <0.01; ***, *p*-value <0.001, ****, *p*-value <0.0001, Mann-Whitney U test with the Benjamini-Hochberg correction for false discovery rate.

Importantly, addition of increasing concentrations of 19F CFS (10%, 20%, 40% (v/v)) to 19F biofilms did not reduce cell counts of 19F 24h post inoculation, suggesting that the 19F strain is immune to its own secreted inhibitory molecules (**Figure S1**).

These results support the hypothesis that secreted molecules of proteinaceous nature are involved in the inhibitory activity of 19F strain, not excluding the involvement of additional mechanisms.

### The genome of strain 19F encodes several bacteriocin-associated clusters

Pneumococcal genomes have been described to encode a large set of bacteriocin gene clusters (6, 12, 24). Together with the results obtained with the CFS of the 19F strain, this prompted us to conduct an *in silico* search for bacteriocin-associated gene clusters in the annotated genome of strain 19F. We found seven loci that could be grouped into three categories according to the classification adopted by Javan et al. (12): bacteriocin-like peptide (Blp) bacteriocins, streptolancidins, and streptococcins (**Figure 2**). Among the Blp bacteriocins, we identified both the ubiquitous *blp* locus containing the regulatory genes, transporters and bacteriocin-immunity region, as well as a duplication of the *blpK* bacteriocin gene and its cognate immunity gene *pncF*, located upstream of the *comAB* genes, both known to be involved in competition (11). The remaining loci identified were those encoding for streptolancidin D, streptococcins A and C, and incomplete versions of loci encoding streptococcins B and E, both lacking the bacteriocin-encoding gene. Of note, apart from the complete version of the streptococcin B encoding locus, none of the latter is known to promote intraspecies competition (9).

**Figure 2.**
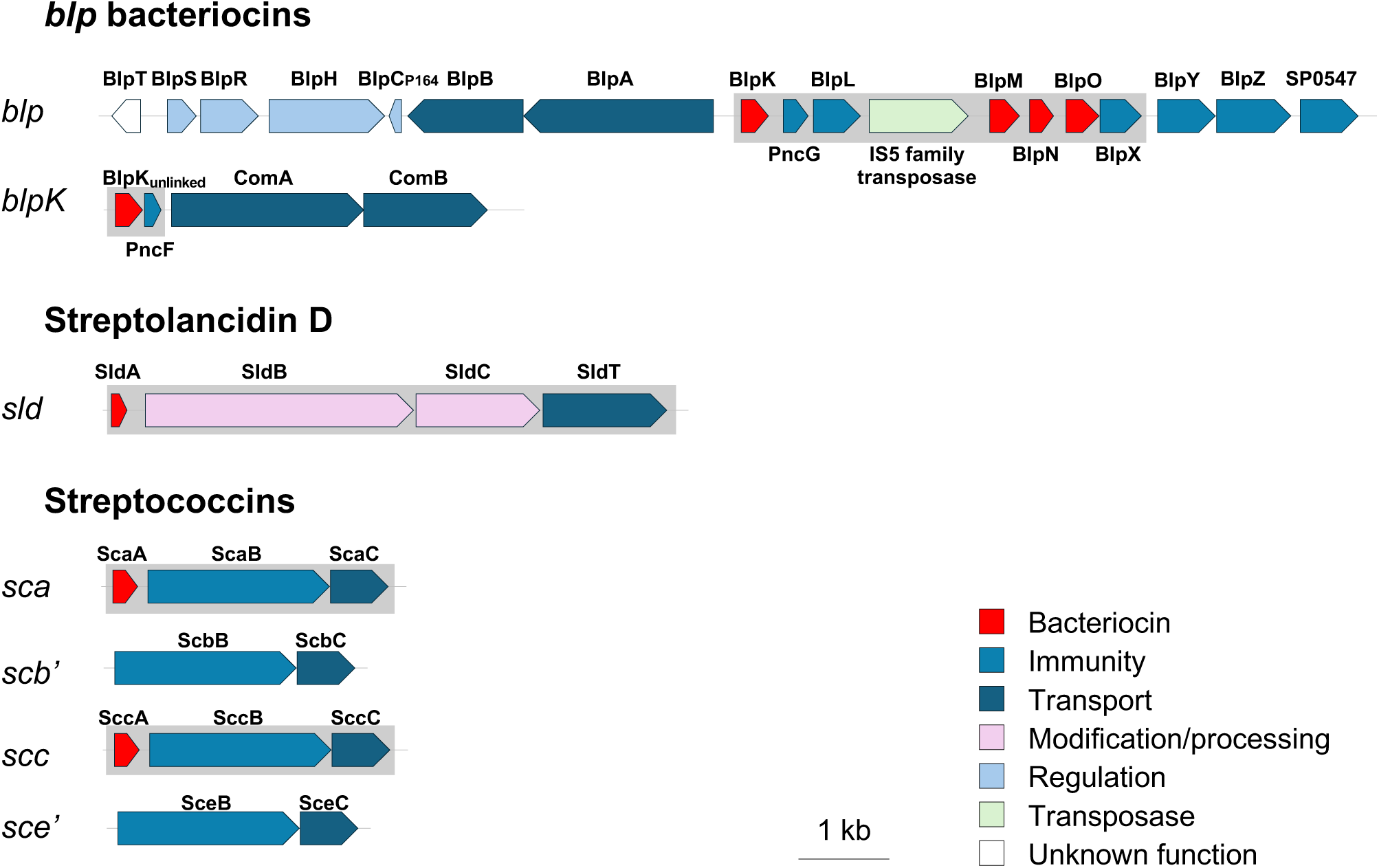
The genome of strain 19F encodes multiple bacteriocin loci. A schematic representation of gene clusters is shown. Genes are colored by function: putative bacteriocins in red, putative immunity proteins in blue, transporters and associated proteins in dark blue, modification proteins in pink, regulators in light blue, and genes of unknown function in white. Deletion-mutant strains were constructed for bacteriocin-containing regions, indicated by light gray boxes. This was done for all but *scb*’ and *sce*’ loci, as these loci are incomplete, do not contain the bacteriocin-encoding gene and, thus, are not predicted to be associated with inhibitory activity. The genome of 1990-19F has been deposited in the NCBI database under accession number SAMN08291405.

Coupled with the differences in bacteriocin-content identified in the genome of NT strain (**Figure S2**), these findings show that strain 19F contains an extensive repertoire of bacteriocins that could drive its inhibitory activity towards the NT strain.

### Secreted streptolancidin D from strain 19F inhibits NT biofilms

To test the involvement of each bacteriocin loci in the inhibitory activity of strain 19F, deletion mutants of each locus were generated individually (**Figure 2**). This was done by deleting the entire locus or just its bacteriocin-immunity region. This strategy was applied to all loci identified, except for the streptococcin-encoding loci lacking the bacteriocin gene, as these are not expected to promote inhibition.

To test for activity, CFS of early stationary-phase planktonic cultures of each mutant were obtained in CDM and added to biofilms of NT strain at the time of biofilm inoculation in a final concentration of 20% (v/v). The CFS of mutant 19FΔ*sldA-T* displayed significantly lower inhibitory activity compared to that of the 19F wild type strain, while the CFS of mutants for the Blp bacteriocins (19FΔBIR and 19FΔ*blpK*) showed comparable inhibitory activity. In contrast, CFS of mutants for streptococcin bacteriocins (19FΔScc and 19FΔSca) showed greater inhibitory activity than CFS from the wild type strain (**Figure 3A**).

**Figure 3.**
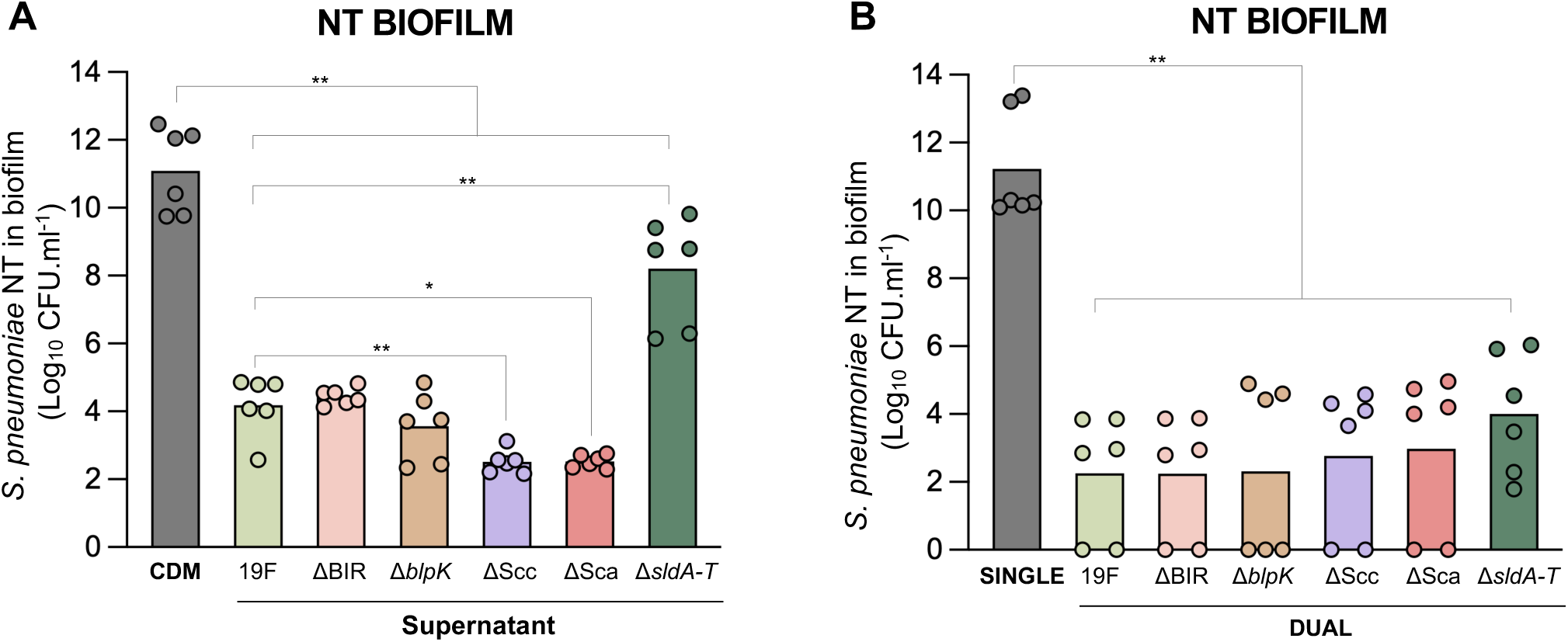
**(A) The supernatant of strain 19FΔsldA-T displays reduced inhibitory activity against strain NT.** Biofilms were grown in CDM for 24h and 20% (v/v) of 19F supernatants (wild type and mutants) obtained from early-stationary phase cultures were added at time of inoculation. CDM, no addition of supernatant, where CDM was added in the same proportion as supernatants; CFU.mL^-1^ were counted after inoculation of serial dilutions of NT biofilms. **(B) Bacteriocin-deficient mutants display inhibitory activity against NT strain comparable to the 19F wild type strain in dual-strain biofilms.** Bacteriocin-deficient mutants were tested in single- and dual-strain biofilms with the NT strain. The graphic is showing NT CFU.mL^-1^ of the NT strain when in co-culture with each 19F variant after 24h biofilm growth in CDM. Black bars indicate geometric means. Graphics represent data from three independent experiments, each with intra-experiment duplicates. *, *p*-value <0.05; **, *p*-value <0.01, Mann-Whitney U test with the Benjamini-Hochberg correction for false discovery rate.

In addition to results described, this experiment showed that while deletion of the *sldA-T* locus dramatically decreases inhibition by 19F, it is not sufficient to abolish the inhibitory activity of the CFS of the strain. This is in agreement with what was observed for the protease-treated supernatant (**Figure 1**), suggesting at least one additional mechanism of 19F-mediated inhibition.

This was further confirmed by testing the inhibitory activity of each mutant against the NT strain in dual-strain biofilms (**Figure 3B**). In an experimental setting in which each 19F variant is in the same biofilm with the NT strain, all mutants inhibited the NT strain to levels comparable to those of the wild type 19F. These dissimilar results obtained with the two strategies used to test mutant activity highlight that intraspecies competition is highly complex and there are likely several mechanisms cooperating for the strong inhibition observed.

In summary, we demonstrate that there are multiple mechanisms driving the 19F-mediated inhibition of strain NT and that products from *sldA-T* are major contributors to inhibition in a contact-independent manner.

### Streptolancidin D promotes competition *in vitro* and *in vivo* and mediates intra-species growth inhibition

To explore the inhibitory role of the *sldA-T* locus, we constructed congenic variants in the background of strain 19F: 19F_GFP (wild type, streptolancidin D producer, GFP-labelled), 19F’Δ*sldA-T* (streptolancidin D deficient, non-labelled); a streptolancidin D-complemented strain 19F’Δ*sldA-T*::*p3-sldA-T* was also constructed (streptolancidin D producer, non-labelled).

The 19F strain was grown in single and dual-strain biofilms with its non-producer variant and with the complemented strain. After 24h, we observed that the 19F wild type strain inhibited the streptolancidin D deficient strain but not the complemented one, clearly demonstrating its inhibitory role (**Figure 4**).

**Figure 4.**
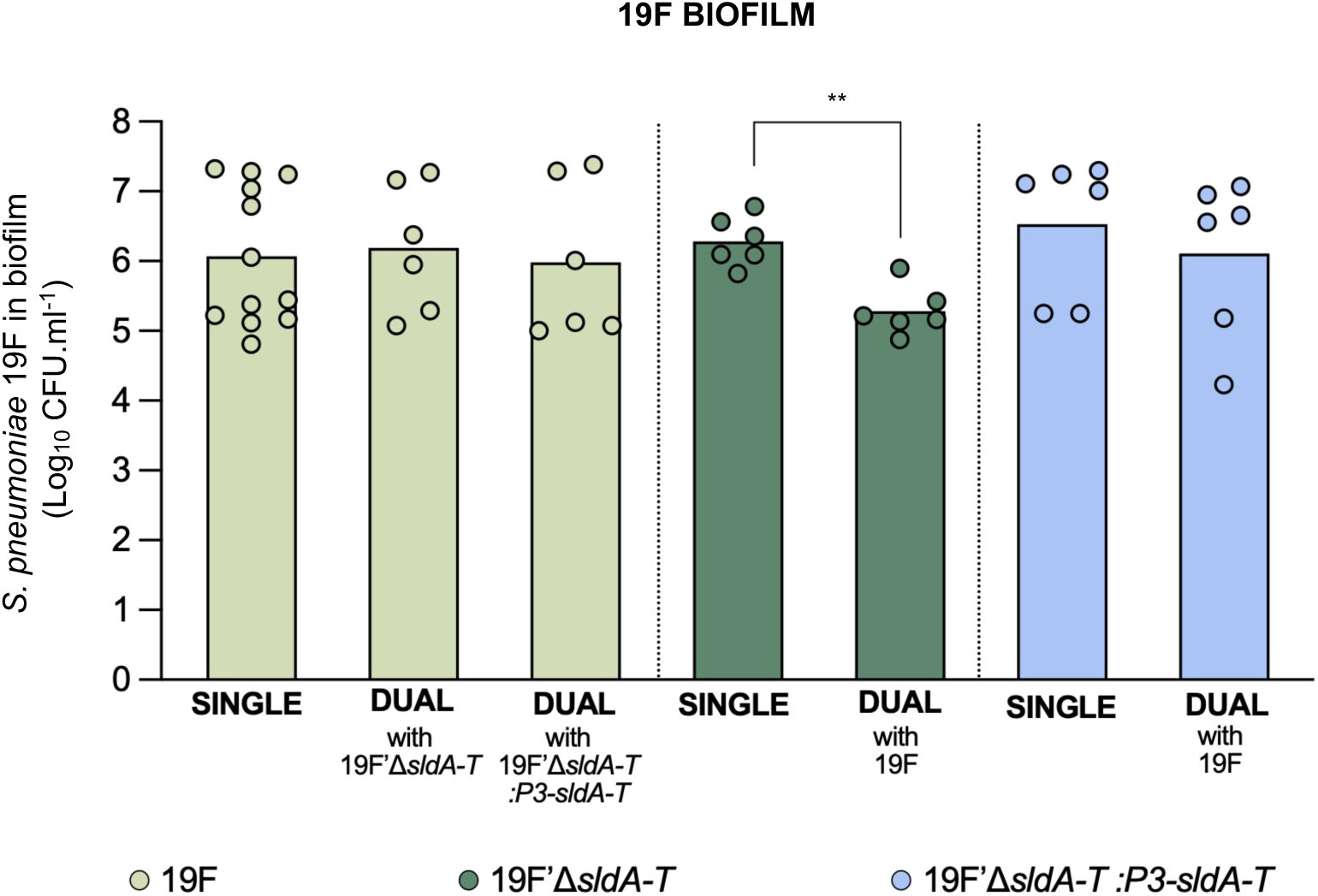
19F inhibits its Streptolancidin D deficient variant but not the complemented strain in dual-strain biofilms. Single- and dual-strain biofilms were grown in CDM for 24h. Co-cultures were prepared in a 1:1 cell ratio with a GFP-labeled (19F, light green) strain and non-labeled (19F’Δ*sldA-T*, dark green or 19F’*ΔsldA-T::P3-sldA-T*, blue) strain. Events.mL-1 were calculated based on cell counts from flow cytometry. Intra-species interactions were assessed by comparison of cell counts of each strain in single and dual-strain biofilms. Plot represents the results of three independent experiments, each with intra-experiment duplicates. Black bars indicate geometric means. **, *p*-value <0.01, Mann-Whitney U test with the Benjamini-Hochberg correction for false discovery rate.

To evaluate the potential of *sldA-T* to promote competition *in vivo*, we used a 7-day co-colonization model. Mice were co-inoculated with strains 19F_GFP (streptolancidin D producer, GFP labelled, chloramphenicol resistant) and 19F’Δ*sldA-T* (streptolancidin D deficient, non-labelled, kanamycin resistant) in a 1:1 ratio. As controls, single colonization experiments with each strain were also conducted. When inoculated independently, both strains colonized the mice for the seven days, reaching similar colonization densities at day 7 post inoculation (**Figure 5A**). In co-colonized mice, the wild type 19F strain (streptolancidin D producer) reached significantly higher nasopharyngeal densities than the non-producer strain 19F’Δ*sldA-T*. Accordingly the competitive index between the two strains decreased from day 0 (1.03 ± 0.087, mean ± SD) to day 7 post-inoculation (0.69 ± 0.076, mean ± SD) (**Figure 5B**).

**Figure 5.**
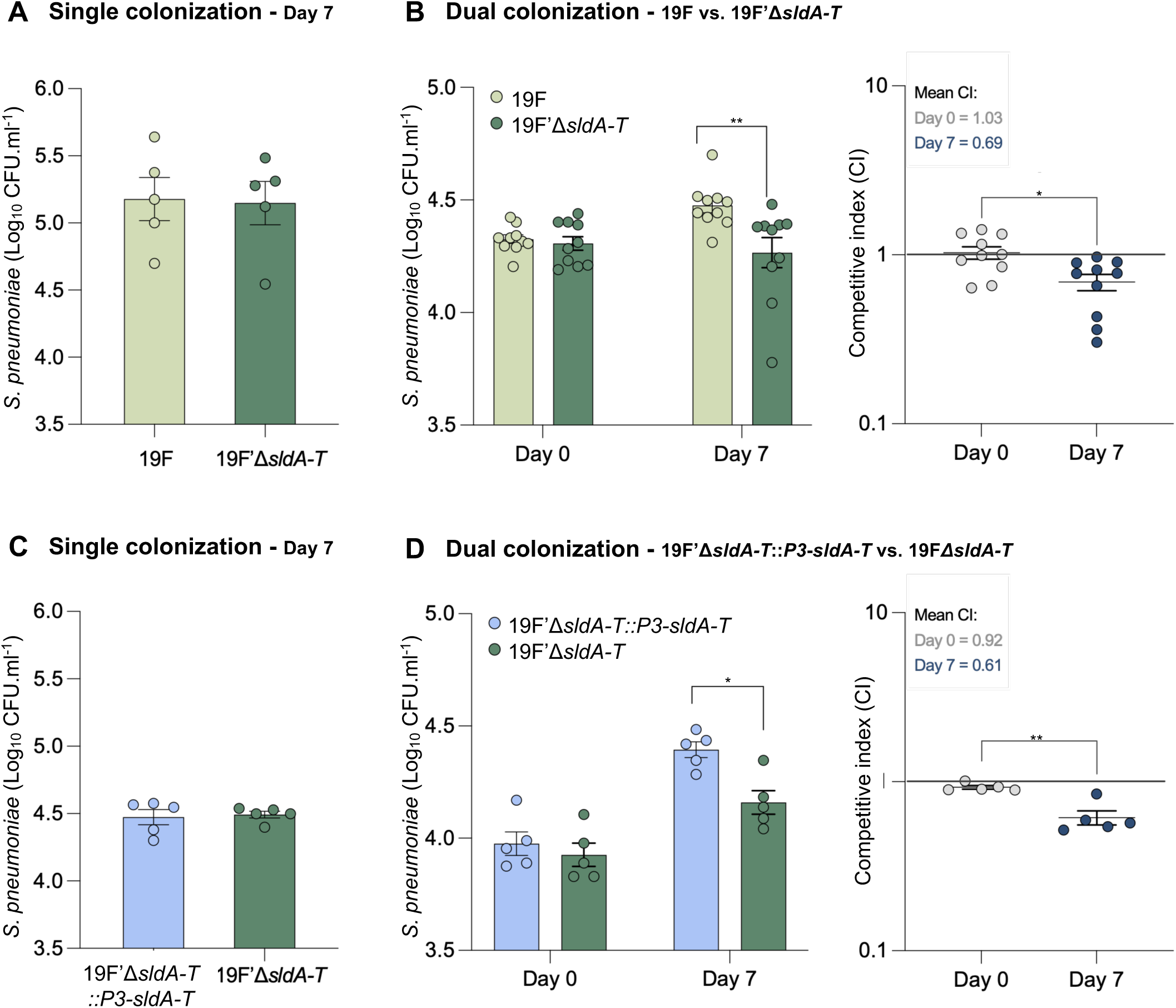
19F streptolancidin D-producing variants outcompete a non-producing variant *in vivo*. Competition experiments were performed using a model of co-colonization in CD1 mice. Mice were intranasally inoculated with 7×10^5^ CFU.mL^-1^, either independently (A) and (C) or in a 1:1 mixture of both strains (B) and (D). Colonization was established for 7 days. Nasal lavages were performed at days 0 and 7 in groups of at least 5 mice. Nasal lavages were serially diluted and plated onto blood-agar plates supplemented with chloramphenicol (to select wild type 19F), kanamycin (to select 19F’Δ*sldA-T*) or kanamycin and spectinomycin (to select 19F’Δ*sldA-T*::*P3-*Δ*sldA-T).* The competitive index was calculated by dividing the CFU ratio 19F::19F’Δ*sldA-T* recovered from the colonized tissue (output ratio) by the CFU ratio of 19F::19F’Δ*sldA-T* present in the inoculum (input ratio) (B) or by dividing the CFU ratio 19F’Δ*sldA-T:P3-sldA-T*::19F’Δ*sldA-T* recovered from the colonized tissue (output ratio) by the CFU ratio of 19F’Δ*sldA-T*::*P3-sldA-T*::19F’Δ*sldA-T* present in the inoculum (input ratio)(C). Black bars indicate geometric means and standard error of the mean. **, *p*-value <0.01, 2-way ANOVA for colonization densities, Mann-Whitney U test for competitive indices.

To further test whether this competitive advantage was specifically associated with streptolancidin D production, an additional co-colonization experiment was performed using the 19F’Δ*sldA-T* mutant and the complemented strain 19F’Δ*sldA-T*::*p3-sldA-T*. Similar to the previous experiment, both strains colonized mice independently at similar levels (**Figure 5C**). Accordingly, in co-colonization experiments, the streptolancidin D-producing complemented strain outcompeted the non-producer mutant during co-colonization, reaching significantly higher nasopharyngeal densities at day 7 post-inoculation (competitive indices of 0.92 ± 0.049 and 0.61 ± 0.132, mean ± SD, on days 0 and 7 post-inoculation, respectively) (**Figure 5D**). These results further support the role of the *sldA-T* locus in mediating pneumococcal fitness advantage *in vivo*.

Overall, these results confirm the role of the *sldA-T* locus as an active bacteriocin locus that promotes competition in pneumococci, both *in vitro* and *in vivo*.

To determine whether the inhibitory effect associated with streptolancidin D production extended beyond interactions between isogenic variants, cell-free supernatants (CFS) from from streptolancidin D-producing and non-producing strains (19FWT and 19FΔ*sldA-T*, respectively) were tested against 36 *S. pneumoniae* strains isolated from human nasopharyngeal samples. This collection included both vaccine and non-vaccine serotypes, in a total of 36, providing a broad representation of pneumococcal diversity (**Table S4**). Exposure to cell-free supernatant from the streptolancidin D-producing strain (19FWT) resulted in growth inhibition of 12 of the 36 strains tested, whereas CFS from the non-producing strain (19FΔ*sldA-T*) had no inhibitory effect towards any of the strains (**Figure 6 and Figure S3**).

**Figure 6.**
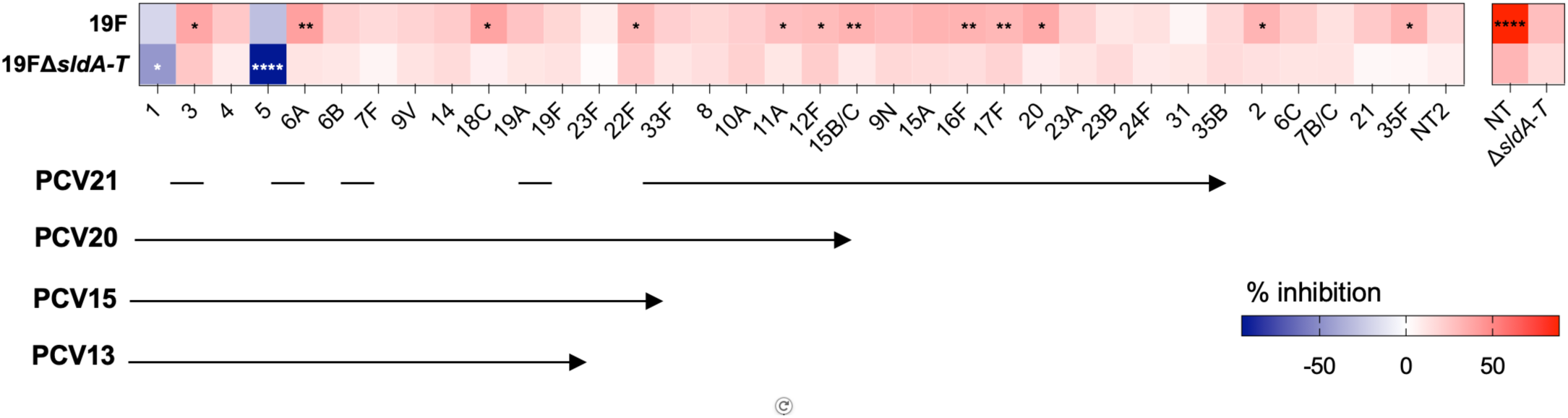
Streptolancidin D - dependent growth inhibition varies across multiple strains. Epidemiologically relevant *S. pneumoniae* strains representing serotypes included in pneumococcal conjugate vaccines (PCV) and non-vaccine types were grown CDM in the presence and absence of cell-free supernatants of WT or Δ*sldA-T* strains. The heatmap represents the percentage of growth inhibition across strains grown for 24 h. Percentage inhibition was calculated from the area under the growth curve (AUC) relative to control (see Figure S3). Positive values (red) indicate growth inhibition, while negative values (blue) indicate growth promotion. Lines underlying serotypes indicate serotypes included in the corresponding pneumococcal conjugate vaccine (PCV). Colours represent the magnitude of the effect. Asterisks indicate statistically significant differences relative to control conditions. Heatmap represents data from three independent experiments, each with intra-experiment duplicates. *, *p*-value <0.05; **, *p*-value <0.01; ****, *p*-value <0.001, one-way ANOVA with Tukey’s with multiple comparisons test.

These findings demonstrate that the inhibitory activity of streptolancidin D, although strain dependent, is not restricted to isogenic strain pairs but can also affect genetically distinct pneumococcal strains.

### Streptolancidin D is distributed across phylogenetically diverse pneumococcal lineages and has a conserved genomic location

To determine the prevalence and phylogenetic distribution of streptolancidin D, we screened a collection of 7,548 pneumococcal genomes (25) for the presence of the *sldA-T* locus. A total of 885 genomes were found to carry the locus, representing 11.7% of the entire dataset. To contextualize the phylogenetic distribution of the *sldA-T* locus, we constructed a maximum-likelihood phylogenetic tree based on core genome alignments of all multilocus sequence types (STs) in which the *sldA-T* locus was detected in at least one representative strain (n=1,194 genomes, **Table S1**). The tree was annotated with *sldA-T* presence/absence, serotype, sequence type, and Global Pneumococcal Sequence Cluster (GPSC) designation (**Figure 7A**).

**Figure 7.**
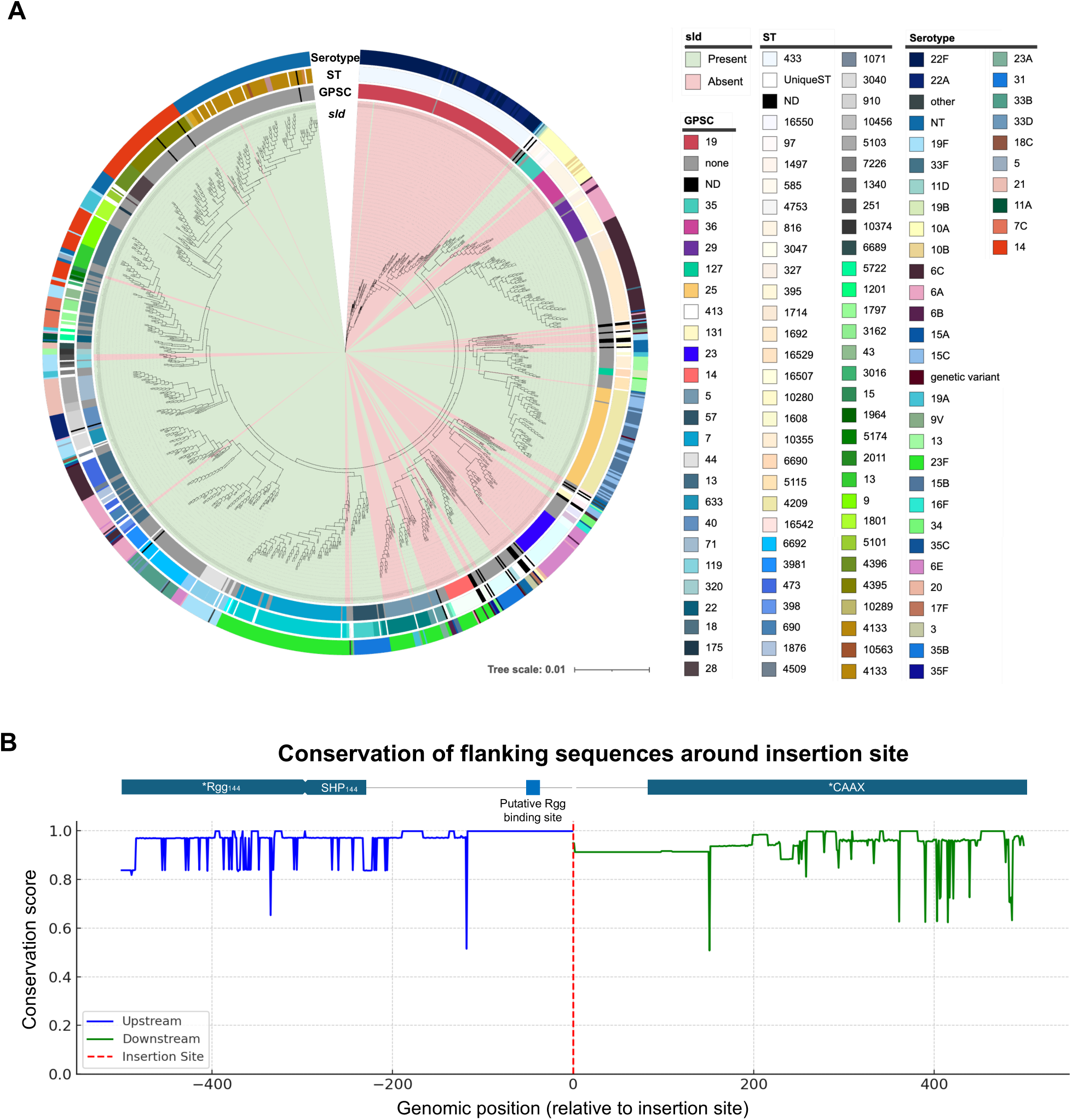
**(A) Distribution of the *sldA-T* locus across genetic backgrounds defined by multilocus sequence type (ST).** The figure represents a maximum-likelihood phylogenetic tree based on core genome alignments of all sequence types (STs) in which the *sldA-T* locus was detected in at least one representative strain (n=1,194 genomes). Genomes were annotated using Prokka (4), core-gene alignments were performed with Roary (9) and the phylogenetic tree was constructed with FastTree (11). Metadata (*sldA-T* presence or absence, GPSC type, MLST type and serotype) were added using the iTOL tool (12). *sldA-T* presence or absence was determined with BLAST. Tree scale indicates number of nucleotide substitutions per site. **(B) Conservation of upstream and downstream sequences flanking the insertion site of a genetic locus across bacterial genomes.** The plot shows conservation scores for the last 500 bp upstream (blue) and first 500 bp downstream (green) of the insertion site. Conservation at each position represents the proportion of aligned sequences that share the most frequent nucleotide. The insertion site is indicated by the red dashed line at position 0. The high degree of sequence conservation in both flanks (average identity: 92.6% upstream, 93.5% downstream) supports the hypothesis that the locus inserts at a consistent genomic location across all genomes analyzed. Plot is showing trimmed conservation of flanking sequences after alignment. Annotation of flanking regions is shown above plot, where asterisks indicate interrupted genes by the 500 bp criterium.

The *sldA-T* locus was detected in 157 STs, from which 74 corresponded to STs with a single isolate in the collection and were classified as Unique STs (**Figure 7A**). The genomes encoding the *sldA-T* locus belong to 31 GPSCs, although approximately half of the genomes in which the locus was found did not have an attributed GPSC at the time of the analysis.

Among the total genome collection, the *sldA-T* locus was most common in STs 311 and 4209, associated with GPSCs 7 and 25, respectively. Overall, most isolates within a given ST carried the locus (**Figure 7A**), while other STs consistently lacked it (**Table S2**) or it was carried by a small number of isolates (e.g., STs 13, 90 and 433) indicating background-specific presence or absence of the *sldA-T* locus. (**Figure 7A, Table S2**). Some STs (e.g., STs 172 and 242) showed variable presence of *sldA-T*, suggesting horizontal gene transfer or differential loss.

With regards to serotype, the locus was more frequently found in genomes of non-encapsulated strains and in strains of serotypes 23F, 14, 6C, 15B/C, 6A, and 19F (serotypes targeted by one or more of the currently licensed pneumococcal conjugate vaccines) (**Table S3**). Importantly, within each serotype, only a subset of the genomes carried the *sldA-T* locus, suggesting that genetic background or recombination play a larger role than serotype alone in determining the presence of this locus.

To assess whether *sldA-T* has a conserved genomic location, we aligned the 500 bp regions upstream and downstream of its insertion site across the bacterial genomes that contained the locus (n=885). The resulting alignments showed high sequence conservation, with an average identity of 92.6% in the upstream region and 93.5% in the downstream region. A plot of the per-position conservation scores further confirmed the consistency of these flanking sequences, with a clear transition at the insertion site. These results strongly support the hypothesis of site-specific insertion of the locus into a conserved genomic region (**Figure 7B**).

Taken together, these results suggest that while *sldA-T* is not ubiquitous, its presence across diverse phylogenetic backgrounds and conserved insertional context point to a potentially important, lineage-associated role in pneumococcal biology.

### Streptolancidin D production is regulated by the Rgg_144_/SHP_144_ quorum sensing system

Considering the conservation of the genomic context of *sldA-T* across pneumococcal genomes, we explored the regulation of the locus. The *sldA-T* locus is encoded downstream of the Rgg_144_/SHP_144_ quorum sensing system, previously shown to regulate downstream genes involved in environmental adaptation, namely biofilm formation and virulence in a PMEN-1 lineage strain and in the reference strain D39, respectively (21, 22). We aligned the *sldA-T* promoter region of strain 19F with the Rgg_144-_regulated promoters in these strains and observed very high conservation (**Figure S4**).

Thus, we investigated whether the Rgg_144_/SHP_144_ system controls streptolancidin D production in strain 19F.

To this end, we constructed luciferase reporters in the background of the 19F strain by insertion of the firefly luciferase gene downstream of the promoter region of *sldA-T*, with and without *shp144*, generating strains 19F’::P*_sld_*-luc and 19F’Δ*shp*::P*_sld_*-luc. These reporter strains were grown for 18h in CDMgal supplemented with luciferin in the presence and absence of exogenously added synthetic 19FSHP C12 (1µM, NZYTech, **Table S1**). We selected the C-terminal 12 amino acids because previous work has shown this constitutes the active SHP_144_ peptide (20, 22, 25).

Results showed high levels (maximum RLU/OD of 1.4×10^7^) of P*_sld_* promoter activity in the wild type strain, regardless of the presence of exogenously added cognate 19FSHP C12, suggesting that secretion of the native peptide is sufficient for maximum promoter activity. On the other hand, in the 19F’Δ*shp*::P*_sld_*-luc, P*_sld_* activity was only detected when exogenous 19FSHP C12 was added (maximum RLU/OD of 1.1×10^6^), showing the Rgg_144_/SHP_144_ drives *sldA-T* expression (**Figure 8**). Of note, addition of synthetic 19FSHP C12 at higher concentration (10 µM), significantly hampered growth of both reporter strains and could not induce activation of P*_sld_* to levels expressed by the wild type strain, even at higher a concentration of inducer (**Figure S5**).

**Figure 8.**
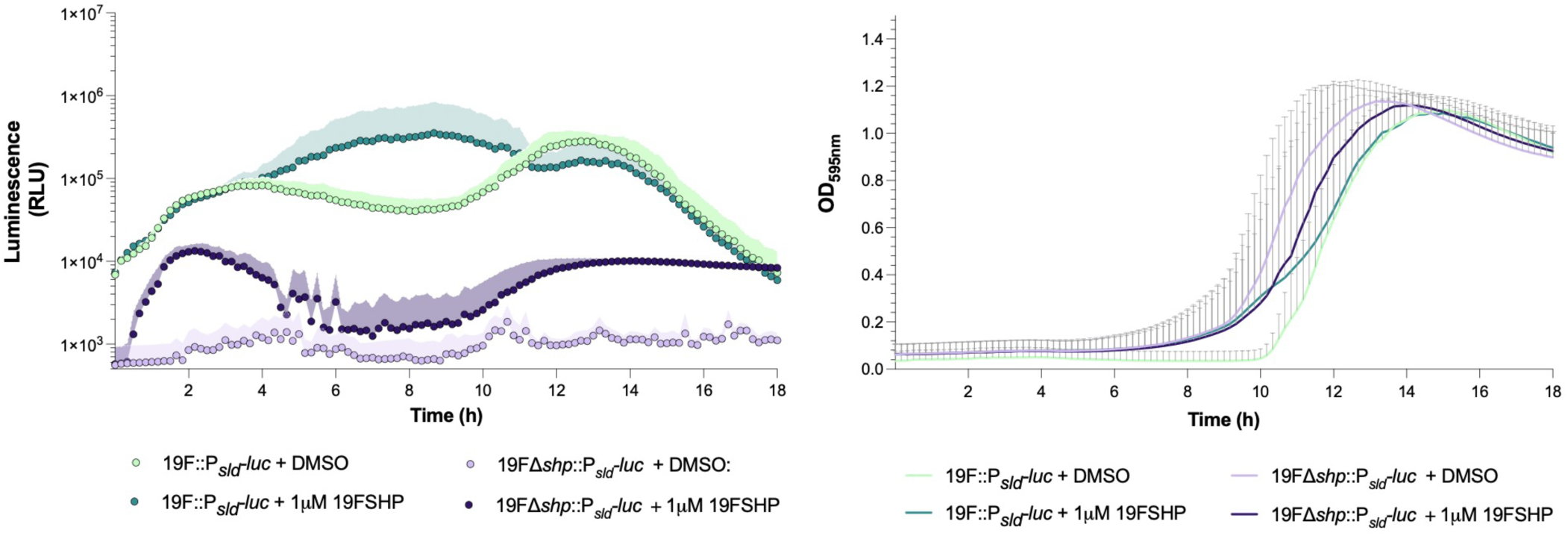
P*_sldA-T_* promoter is activated by SHP_144_. Reporter strains containing the luciferase gene upstream of the ATG of *sldA* gene were constructed in the background of strains 19F’ (19F’::P*_sld_-luc*, green) and 19F’Δ*shp* (19F’Δ*shp:*:P*_sld_-luc*, purple). Growth curves were performed in the presence (dark colour) and absence (light colour) of 1μM of synthetic 19FSHP C12. Luminescence (left) and optical density (OD_595nm_) (right) were measured at 10 min intervals and plotted. Dots indicate luminescence, lines represent growth curves. Shaded areas and bars represent standard deviation of the mean luminescence and optical density, respectively. Plotted are results of three independent experiments, each with intra-experiment duplicates.

These results demonstrate that streptolancidin D production is regulated by the Rgg_144_/SHP_144_ system, leveraging a conserved genomic context for coordinated control.

### Cumulative bacteriocin activity and competence-mediated fratricide drive intra-species competition

Our results identified streptolancidin D as a major contributor to inhibition. Still, the incomplete loss of activity observed upon deletion of the *sldA-T* locus suggested that additional mechanisms contribute to the inhibitory phenotype of strain 19F. Given that the 19F genome encodes multiple bacteriocin loci, we hypothesized that inhibition could result from the combined activity of these loci. Differences between the bacteriocin repertoires of strains 19F and NT (the *blp* locus bacteriocin content and the absence of Streptococcin A in NT, Figure 2 and Figure S2) supported this hypothesis.

To address this, we constructed multiple deletion mutants targeting combinations of bacteriocin loci, generating double (19FΔBIRΔ*sldA-T*), triple (19FΔBIRΔ*sldA-T*Δ*blpK*), and quadruple (19FΔBIRΔ*sldA-T*Δ*blpK*ΔSca) mutants. Simultaneous deletion of bacteriocin-associated loci progressively reduced the inhibitory activity against the NT strain, with the strongest effect observed for the triple deletion mutant (**Figure 9**). In contrast, additional deletion of the Sca locus did not further decrease inhibition, suggesting that this locus is unlikely to contribute substantially to NT inhibition under these conditions.

**Figure 9.**
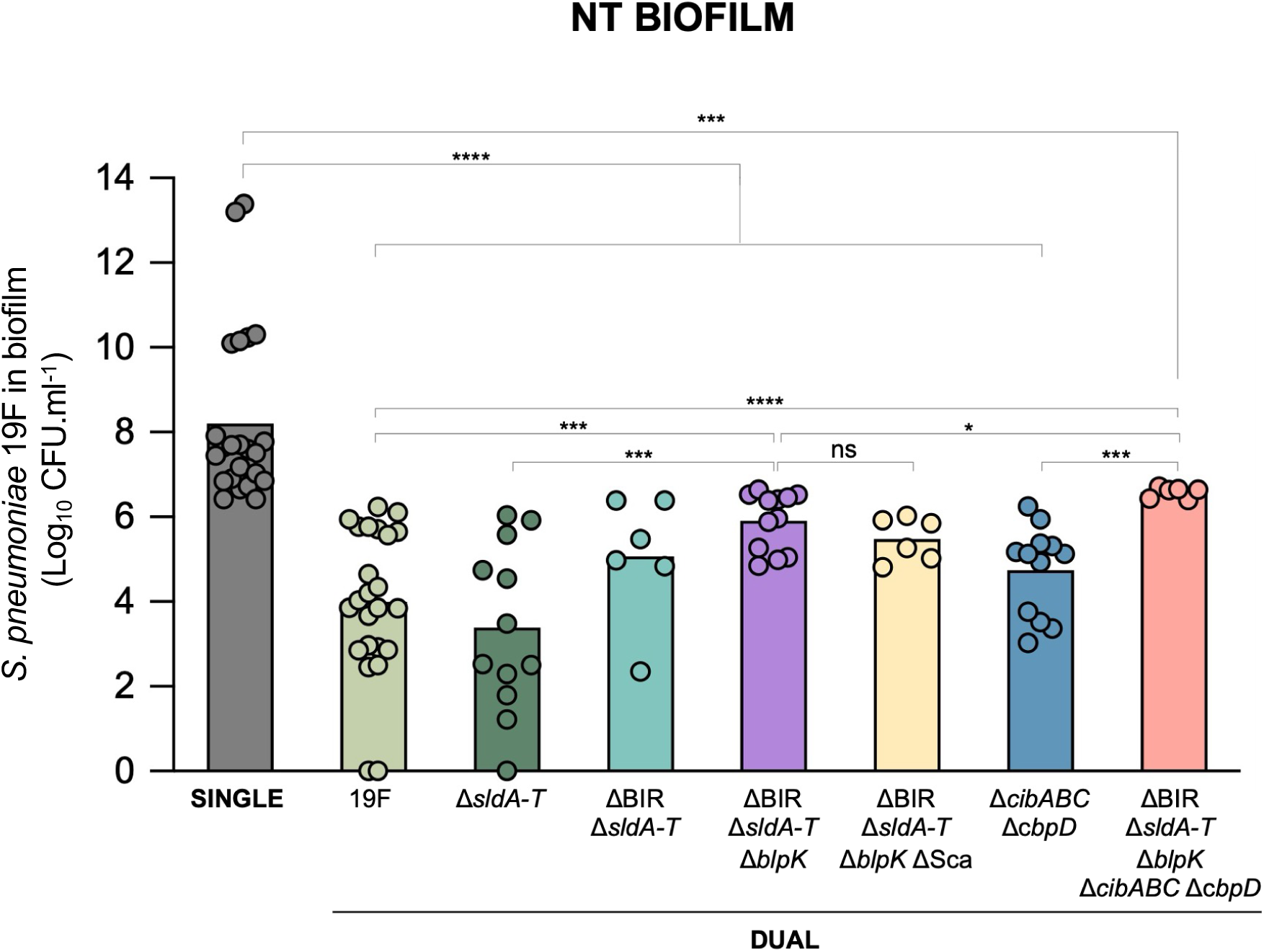
Cumulative inhibitory activity of bacteriocins and competence-mediated fratricide drive intra-species competition. Bacteriocin- and fratricide-deficient mutants were tested in single- and dual-strain biofilms with the NT strain. The graphic shows CFU.mL^-1^ of the NT strain when in co-culture with each 19F variant after 24h biofilm growth in CDM. Black bars indicate geometric means. Graphics represent data from at least three independent experiments, each with intra-experiment duplicates. *, *p*-value <0.05; ***, *p*-value <0.001, ****, *p*-value <0.0001, Mann-Whitney U test with the Benjamini-Hochberg correction for false discovery rate.

Importantly, even combined deletion of major bacteriocin loci did not fully restore NT growth, indicating that bacteriocins alone do not fully account for the observed phenotype. This observation, together with the discrepancy between contact-dependent and contact-independent assays (**Figure 3**), pointed to the involvement of an additional mechanism requiring direct cell-cell interaction, being competence-mediated fratricide a likely candidate. To test this hypothesis, the fratricide effectors *cibABC* and *cbpD* were deleted in the background of the 19F wild-type strain (19FΔ*cibABC*Δ*cbpD*) and in the triple mutant background (19FΔBIRΔ*sldA-T*Δ*blpK*Δ*cibABC*Δ*cbpD*). The triple mutant background was selected based on its stronger reduction in inhibitory activity compared to the quadruple mutant. This was consistent with the observation that the single Sca deletion mutant exhibited increased inhibitory activity against the NT strain (**Figure 3**). The combined deletion of bacteriocin loci and fratricide effectors resulted in the lowest inhibitory activity against the NT strain, supporting the contribution of contact-dependent mechanisms to the 19F inhibitory phenotype (**Figure 9**).

Together, these results demonstrate that intra-species competition mediated by strain 19F is a multifactorial process, in which Rgg_144_/SHP_144_-controlled streptolancidin D acts as a key component within a broader network of bacteriocins and competence-associated mechanisms that collectively determine competitive outcomes.

## DISCUSSION

In this study, we identified and characterized streptolancidin D, a previously uncharacterized bacteriocin encoded by the *sldA-T* locus, and demonstrated its role in intra-species competition in *Streptococcus pneumoniae.* Using a naturally colonizing strain (19F) with potent inhibitory activity against other pneumococci, we showed that *sldA-T* contributes significantly to bacterial antagonism both *in vitro* and *in vivo*.

Streptolancidin D, encoded by *sldA*, is part of an operon containing three additional genes: *sldB*, *sldC* and *sldT*, predicted to encode a dehydratase, a cyclase and a lantibiotic transporter (12). No dedicated immunity genes are present in the operon. Still, the lack of inhibition of the wild type 19F towards the *sldA-T* complemented strain suggests that the immunity mechanism may be encoded within the operon. One hypothesis could be a dual role for the SldT transporter, as being involved in export of the bacteriocin and also having an immunity role, as described for other lantibiotics, such as Nisin and Lacticin 3147 (26, 27).

Streptolancidin D plays a significant role in enhancing pneumococcal competitive fitness, particularly in structured environments such as biofilms and during co-colonization *in vivo*. We further demonstrate that the inhibitory activity of streptolancidin D is not restricted to interactions between isogenic variants. Supernatants from a streptolancidin D-producing strain inhibited several pneumococcal strains of diverse vaccine and non-vaccine serotypes, whereas that from a non-producing variant had no detectable effect. Although susceptibility varied between strains, these findings indicate that streptolancidin D mediates interactions among genetically distinct pneumococci, with variability in sensitivity likely reflecting differences in resistance and/or immunity mechanisms.

Deletion of *sldA-T* resulted in a marked reduction in inhibitory activity in supernatant-based assays, highlighting its contribution to secreted antimicrobial function. However, this deletion did not abolish inhibition in dual-strain biofilms, suggesting that additional mechanisms, such as other bacteriocins or non-proteinaceous factors, may act redundantly or synergistically to sustain competitive pressure. In fact, the progressive reduction in inhibitory activity observed following deletion of multiple bacteriocin loci indicates that competition mediated by strain 19F relies on the cumulative action of several antimicrobial systems rather than on streptolancidin D alone. Furthermore, deletion of the fratricide effectors CibABC and CbpD in a bacteriocin-deficient background resulted in an additional reduction in inhibition, supporting a contribution of competence-mediated killing. Together, these observations reveal that pneumococcal competition is a multifactorial process involving both secreted bacteriocins and contact-dependent mechanisms. Such redundancy may increase the robustness of competitive interactions in the dynamic and densely populated nasopharyngeal niche. In structured environments, factors such as cell-cell contact, bacteriocin cooperation, local peptide accumulation, and extracellular matrix composition are known to influence the outcome of inter-strain interactions (7, 15, 28, 29).

Our comparative genomics analysis shows that the *sldA-T* locus is consistently encoded downstream of the Rgg_144_/SHP_144_ quorum sensing system, maintaining a tight regulatory architecture across genomes, aligning with the description of Javan et al (12). The consistent insertion of the *sldA-T* locus downstream of the Rgg_144_/SHP_144_ genes across pneumococcal genomes suggests selective pressure to maintain this regulatory architecture. This positioning is reminiscent of other bacteriocin systems in *S. pneumoniae*, where a signaling peptide controls expression of antimicrobial genes in a cell density-dependent manner (6, 14, 30). Quorum sensing-regulated antimicrobials allow bacteria to optimize effort, target competitors effectively, and synchronize population behaviors, giving them a strong ecological advantage in structured, resource-limited environments like the nasopharynx.

Structure-function studies have confirmed that SHP_144_ plays a critical role in Rgg_144_ activation and subsequent transcriptional regulation, impacting nutrient utilization and stress response pathways in pneumococcus (22, 25). Cuevas et al. have also demonstrated that Rgg_144_ activates transcription of the *vp1* operon (syntenic to *sldA-T*) in the presence of SHP_144_. Functional studies confirmed that deletion of *rgg144* or *shp144* significantly reduces *vp1* expression, leading to diminished biofilm formation and attenuated virulence in mouse models (21). In our study, the promoter activity of *sldA-T* was abolished in the Δ*shp* mutant and partially restored by addition of the exogenous peptide, supporting direct regulation by this quorum sensing system. The inability of synthetic SHP_144_ to fully recapitulate wild-type expression levels suggests that native SHP_144_ secretion and processing are finely tuned and potentially optimized for strain-specific regulatory dynamics.

Our analyses revealed that streptolancidin D is broadly but unevenly distributed across the pneumococcal population, occurring in multiple sequence types and GPSCs, yet often restricted to specific genetic backgrounds. While certain serotypes showed higher prevalence of the locus, its presence seemed to be more tightly linked to genomic background than to capsule type alone. In line with this, we have observed that *sldA-T* is present in both vaccine and non-vaccine serotypes without evidence of differential prevalence between these groups. This supports the view that the presence of this locus and its maintenance in the population is likely driven by lineage-specific evolutionary dynamics. Streptolancidin D could serve as a colonization factor, enabling its carriers to exclude competing pneumococci and maintain niche dominance. This fully aligns with the hypothesis that bacteriocins, beyond mediating competition, may play roles in shaping population structure following PCV implementation (31).

In conclusion, this study expands our understanding of the diversity of antimicrobial strategies in *S. pneumoniae*, highlight the interplay between quorum sensing and competitive behavior, and reinforce the idea that bacteriocins can influence not only microbial interactions but also long-term population structure. Further research exploring the structure, mechanism of action, and immunity determinants of streptolancidin D will be essential for fully understanding its role in pneumococcal ecology and evolution.

## METHODS

### S. pneumoniae strains

The colonization strains used in the study, 1990-19F and 7031-NT, were isolated from nasopharyngeal samples of healthy children (32). Fluorescently labelled variants of these strains were generated and validated in a previous study (23). All other 19F variants generated for the purpose of this study are listed in **Table S4**.

### Biofilm growth conditions

Biofilms were grown in CDMgal as previously described (23). Briefly, 10^5^ bacteria in 2.5mL of CDMgal were seeded in 24-well plates and incubated at 34°C in 5% CO_2_ for 24 hours, except if otherwise indicated. In dual-strain biofilms, strains were mixed in a 1:1 ratio and grown as described above. For comparative analysis, single-strain biofilms of the same strain variants were obtained in parallel. At 24h, biofilms were resuspended in 300µL of PBS and either serially diluted for colony forming units (CFU) counts (NT selection on 1µg/mL erythromycin) or sonicated to separate aggregates prior to flow cytometry analysis, as previously described (23). For all experiments three biological replicates were done and each included at least two technical replicates.

### Supernatant experiments

Cell free supernatants were obtained from early stationary-phase planktonic cultures of strain 1990-19F_GFP in CDMgal. The culture was centrifuged, the pellet was discarded, and supernatants were filtered using a 0.2µm vacuum filter.

To investigate the molecule(s) mediating amensalism, 1990-19F_GFP cell-free supernatants were treated with 1mg/mL proteinase K (1h at 37°C, followed by inactivation at 95°C for 10min) or heated at 65°C for 20min. Biofilms of strain 7031-NT were grown for 24h, as described, in the absence (control) or presence of 20% (v/v) 1990-19F_GFP supernatant under the following conditions: (i) untreated; (ii) proteinase K-treated; and (iii) heated at 65°C. After 24h, biofilms were resuspended in 300µL PBS 1x, sonicated, and cell counts determined by flow cytometry, as previously described(23).

To assess the inhibitory effect of WT bacteriocin-deleted 19F variants on bioflim growth of 7031-NT_RFP, untreated cell free supernatants were added to inocula (20% v/v) prior to biofilm seeding and growth was promoted for 24h, as described above.

To assess the inhibitory effect of streptolancidin D on planktonic growth of representative pneumococcal isolates, strains were grown to an OD_600_=0.5, diluted 1:100 and grown CDM with 1600U/mL catalase in the presence and absence of cell-free supernatant (20% v/v) in 96-well plate (final volume 200µL). Absorbance was measured at 595 nm using the Tecan Infinite 200 pro microplate reader at 37 °C every 30 minutes for 24 hours. The percentage of inhibition was calculated by comparing the area under the curve (AUC) obtained under each experimental condition with the AUC of the control condition (CDM), using the following formula: % inhibition=(AUC_CDM_-AUC_condition_)/AUC_CDM_*100.

### Whole genome sequencing and analysis

Genomic DNA was extracted using the MagnaPure equipment, according to the manufacturer’s instructions, and quantified in a Qubit 2.0 fluorometer (Life Technologies). Library preparation and whole genome sequencing (WGS) were carried out using the Illumina Next Seq platform at the Genomics Facility of Instituto Calouste Gulbenkian (IGC).

Quality control, genome assembly and determination of multilocus sequence type (MLST) was performed using INNUca (v.3.1). Genome annotation was done using Prokka (v.1.13.3) (33).

For identification of bacteriocin-associated gene clusters, the annotated assembled genome was submitted to the tools BAGEL4 (34) and antiSMASH v5.0 (35). The outputs from these tools were manually examined and combined to establish more confident gene clusters. Small ORFs not automatically annotated with the Prokka pipeline were searched using the “find open reading frame” (any start codon) tool from CLC Genomics Workbench (Qiagen) and their putative function was examined by BLASTp search against the NCBI and/or the UniProt databases. The genome of 1990-19F has been deposited in the NCBI database under accession number SAMN08291405.

### Construction of deletion mutants

Markerless deletion of the BIR (bacteriocin and immunity region) of the *blp* locus was performed using a previously described genome-editing system(36). Regions flanking the BIR in strain 1990-19F_GFP were amplified and fused by overlap-extension PCR using primers listed in Table S4. The resulting fragment and purified pORI plasmid were digested with *XbaI* and *EcoRI* (New England Biolabs), purified, and ligated using T4 DNA ligase (Thermo Scientific). The ligation product was ethanol-precipitated and transformed into *Lactococcus lactis* 108 by electroporation, as previously described (36). Transformants were selected on SM17 agar supplemented with 0.5% glucose, erythromycin (5 µg/mL), and X-gal (80 µg/mL), and confirmed by colony PCR. Plasmid DNA from positive clones was extracted and transformed into strain 1990-19F_GFP. Transformants were selected on blood agar containing chloramphenicol (4 µg/mL) and erythromycin (5 µg/mL), and screened on TSA supplemented with catalase (4750 U), chloramphenicol, erythromycin, and X-gal. Correct integration was confirmed by colony PCR. To promote plasmid excision, transformants were grown in C+Y_YB_ and plated on TSA containing catalase and X-gal. White colonies were screened for erythromycin susceptibility and viability by replica plating onto TSA with X-gal, blood agar with erythromycin, and non-selective blood agar plates. Deletions were confirmed by colony PCR.

Deletion of *sca* locus in strain 19FΔBIRΔ*sldA-T*Δ*blpK* was done through allelic replacement with a modified version of the PhunSweet cassette (37). Upstream and downstream flanking regions of the *sca* locus were amplified and fused to the *sacB* gene (from D39::PhunSweet) and an erythromycin resistance cassette (*erm*, from F22). The construct was transformed into 19F19FΔBIRΔ*sldA-T*Δ*blpK* and transformants were selected on TSA containing erythromycin (1µg/mL), generating intermediate strains carrying *erm* and *sacB*. Markerless mutants were subsequently generated by transformation with a construct containing only the fused flanking regions, followed by counterselection on TSA supplemented with 10% sucrose. Mutants were confirmed by colony PCR.

Deletion of the remaining bacteriocin loci and fratricide effector genes was performed using a previously described Cre-lox system (38). Target loci were replaced by kanamycin-, tetracycline-, or spectinomycin-resistance cassettes flanked by lox66 and lox71 sites. Flanking regions of each target locus were amplified from genomic DNA. The *lox66-P3-kanR-lox71* cassette was amplified from pKan, *tet(M)* from strain F22 (10), and *spec* from pPEP1_LGZ_PblpT to generate *lox71-P3-tetR-lox66* and *lox66-spec-lox71* cassettes, respectively (Table S4).

PCR products were purified using the Zymoclean™ Gel DNA Recovery Kit (Zymo Research) and assembled by Gibson Assembly (NEB). Nested PCR was used to amplify the final transformation constructs.

When required, resistance markers were excised by transformation with the temperature-sensitive plasmid pJB01 carrying the *cre* recombinase gene, as previously described (38). Transformants were selected on blood agar supplemented with kanamycin (150µg/mL), tetracycline (1µg/mL), or spectinomycin (200µg/mL), as appropriate, and confirmed by PCR.

For all transformations, 1990-19F variants were grown in C+Y_YB_ medium without shaking at 37°C until an OD_600nm_ of 0.5. Cultures were diluted 1:100 in fresh C+Y_YB_ and grown until an OD_600nm_ of 0.1. At that point, cultures were supplemented with 150ng/mL of DNA and 2µM of CSP2 (Mimotopes, Australia), further incubated for 3h and platted onto blood-agar plates supplemented with the appropriate antibiotic or succrose.

### Construction of *sldA-T* complemented strain

The *sldA-T* locus was integrated into the CEP region (39) of the chromosome of 19FΔ*sldA-T* under the control of the P3 constitutive promoter. The backbone of the replicative plasmid pPEP1_LGZ_PblpT was amplified and bunt-end ligated to the construct *p3-sldA-T* after its phosphorylation. Competent *E. coli* DH10 cells were transformed with the ligation product. Plasmid extraction was performed with the ZR Plasmid Miniprep kit (Zymo Research) following manufacturer’s instructions. For confirmation of the orientation of the *p3-sldA-T* fragment in the plasmid, digestion with *HindIII* was used. Purified plasmid was linearized with *KpnI* and used for transformation of strain 19FΔ*sldA-T*, as described above. Transformants were selected based on spectinomycin resistance and confirmed by colony PCR. Primers, strains and plasmid used are listed in **Table S4.**

### *In vivo* competition experiments

For *in vivo* competition experiments, 8-10-week-old female CD1 outbred mice (bred in house at the Leicester University, UK) were intranasally inoculated with 7×10^5^ CFU of strains to be tested in a 1:1 ratio, in 20μl PBS. At days 0 and 7, mice were anesthetized with 5% (v/v) isoflurane over oxygen and then killed by cervical dislocation (at least 5 mice per group) and nasal washes were collected and plated onto blood-agar plates supplemented with either chloramphenicol (to select 1990-19F_GFP), kanamycin (to select 19F’Δ*sldA-T), or* kanamycin and spectinomycin (to select 19F’Δ*sldA-T::P3-sldA-T*) for CFU counts. For control, CD1 mice were inoculated with 7×10^5^ CFU of each strain alone (n=5 per group) and colonization was allowed to establish until day 7, at which point mice were sacrificed, nasal washes collected and platted for CFU counts as described above.

The competitive index was calculated by dividing the CFU ratio 19F:19F’Δ*sldA-T* recovered from the colonized tissue (output ratio) by the CFU ratio of 19F:19F’Δ*sldA-T* present in the inoculum (input ratio) or by dividing the CFU ratio 19F’Δ*sldA-T::P3-sldA-T* :19F’Δ*sldA-T* recovered from the colonized tissue (output ratio) by the CFU ratio of 19F’Δ*sldA-T::P3-sldA-T* :19F’Δ*sldA-T* present in the inoculum (input ratio). A CI less than one indicates that the streptolancidin D-producing strain outcompeted strain 19F’Δ*sldA-T* during colonization, whereas a CI greater than one indicates the opposite.

All animal experiments were conducted under UK Home Office license PP0757060 in accordance with the Animals (Scientific Procedures) Act 1986 and approved by the University of Leicester Ethics Committee. Mice were housed in individually ventilated cages under controlled conditions and monitored regularly post-infection to minimize suffering.

### Construction of luciferase reporters and promoter activity assays

Strains bearing a translational fusion of the *sldA-T* promoter to the luciferase reporter (19F’::P*_sldA-T_-luc* and 19F’Δ*shp*::P*_sldA-T_-luc*) were constructed in the background of strain 1990-19F. First, insertion of the firefly *luciferase* gene downstream of the promoter region of *sldA-T* was achieved by allelic replacement with a fragment containing the *luciferase* gene fused to the upstream and downstream regions of the ATG of *sldA* gene. The *luciferase* and *cat* genes were amplified from pJB11 (**Table S4**) and transformants were selected for resistance to chloramphenicol. Second, deletion of *shp* from the first reporter was achieved by allelic replacement of *shp* gene with the *lox66*-*P3-KanR-lox7* fragment, as described above. These mutants were selected based on kanamycin resistance. All mutants were confirmed by colony PCR. Primers used are listed in **Table S4.**

To test promoter activation both reporters were grown in CDMgal to an OD_600nm_ of 0.5, at which point the cultures were diluted 1:100 in CDMgal supplemented with 0.45mg/mL luciferin (Abcam), and 1600U.mL^-1^ of catalase (Sigma-Aldrich), transferred to a 96-well plate, and either treated with 1mM of synthetic 19FSHP (NZYTech) (**Table S4**) or with 5mM DMSO SHP solvent. The plate was incubated at 37°C for 18h in a BioTek Neo2 Plate Reader and bacterial growth (OD_595_) and luciferase activity (RLU) were measured every 10 minutes.

### *In silico* characterization of *sldA-T* locus and epidemiological analyses

For functional prediction of the proteins encoded by the *sldA-T* genes, the annotations provided by the AntiSMASH and BAGEL databases were used, and the amino acid sequence of each protein was manually searched in the Pfam database to search for domains predictive of the protein function.

In the genome of the 1990-19F, the *sldA-T* locus locates upstream of the quorum-sensing system encoded by *rgg144/shp144* genes. To evaluate if this location was conserved within the population, we performed BLASTn against a collection of well-curated collection of 7,548 pneumococcal genomes kindly provided by Melissa Jansen van Rensburg and Angela Brueggemann and available from the PubMLST database (25) using the entire locus (*sldA-T*) as query. The *sldA-T* upstream and downstream regions (500 bp) of the positive hits were extracted, concatenated and aligned with MAFFT (40). Conservation scores were calculated and plotted using Matplotlib (41).

For phylogenetic analyses, draft assemblies of the PubMLST genomes were annotated using Prokka v1.13.3 (33). All genomes with sequence types (STs) in which at least one genome was a positive hit for *sldA-T* in the BLAST analysis were selected and Roary v3.10.0 was then used to identify core (present in >99 % of isolates) and accessory genes and to generate a core gene alignment (42). The core genome alignment generated by Roary was used to construct a phylogeny using FastTree Version 2.1.9 SSE3 (43). For phylogenetic tree visualization and addition of metadata (*sldA-T* presence, serotype, MLST type and GPSC), the iTOL tool was used (44).

### Statistical analyses

The two-tailed unpaired Mann-Whitney U test with Benjamini and Hochberg correction for FDR was used for comparisons of *S. pneumoniae* viability in single and dual-strain biofilms and in biofilms grown in the presence of cell-free supernatants. A two-way ANOVA was used for comparison of CFUs recovered from each *S. pneumoniae* strain. The Mann Whitney U test was used to compare competitive indices. The one-way ANOVA with Tukey’s multiple comparisons test was used to compare the AUC of growth curves of pneumococcal strains in the presence and absence of cell free supernatant of streptolancidin D producer and non-producer 19F variants. A *p*-value ≤ 0.05 was considered significant. All statistical analyses were performed using GraphPad Prism v9.0 (GraphPad Software Inc., La Jolla, California, USA).

## ACKNOWLEDGEMENTS

The authors are grateful to Morten Kjos (NMBU), Sérgio R. Filipe (FCT NOVA) and João Borralho (ITQB NOVA) for providing plasmids used in the study, and to João M. Lourenço (FCT NOVA) for providing computational resources and support required for phylogenetic tree construction.

The authors kindly acknowledge the technical support provided by the DBS in Leicester, UK, for the *in vivo* work.

This work was supported by FCT - Fundação para a Ciência e a Tecnologia, I.P., through project PneumoISIS (2022.10980.PTDC), MOSTMICRO-ITQB R&D Unit (doi.org/10.54499/UID/04612/2025, UID/PRR/4612/2025) and LS4FUTURE Associated Laboratory (DOI 10.54499/LA/P/0087/2020). C.V. was supported by a post-doctoral fellowship (SFRH/BPD/115280/2016) from FCT.

## AUTHOR CONTRIBUTIONS

C.V and R.S.-L. contributed to the concept and design of the study. CV, HY, and R.S.-L. contributed with reagents and materials. Data acquisition was performed by CV, SD, ARC and OG. Results were interpreted and discussed by all authors. The manuscript was drafted by CV and SD and critically revised by all authors. All authors approved the final version of the manuscript. CV and SD contributed equally to the study.

